# A genetic screen in enteroendocrine cells reveals mechanisms that control protein sensing and GLP-1 release

**DOI:** 10.64898/2025.11.30.691441

**Authors:** Shenliang Yu, Yoochae Lee, Steven C. Boggess, Nicole R Klein, Noam Teyssier, Martin Kampmann, Zachary A. Knight

## Abstract

Enteroendocrine cells (EECs) are the principal nutrient sensors in the gastrointestinal (GI) tract and release hormones such as glucagon-like-peptide 1 (GLP-1) that modulate GI function and appetite. While some of the molecules involved in nutrient sensing within EECs have been described, there have been no systematic studies to map the relevant genes and pathways. Here, we developed a strategy to perform a high-throughput screen for genes that are required for nutrient-induced activation of EECs, and we applied this to probe mechanisms for sensing dietary protein. We found that all of the genes previously proposed to function as protein sensors in EECs are, collectively, dispensable for protein sensing in an EEC cell line. Instead, a screen of >20,000 sgRNAs identified numerous genes associated with mitochondrial respiration as being necessary for this process. We showed through secondary assays that impairing oxidative phosphorylation (OXPHOS) reduced EEC activation and GLP-1 release in response to nutrients but not in response to a non-nutritive stimulus. On the other hand, boosting OXPHOS increased EEC activation and GLP-1 release. These data reveal that intracellular metabolism within EECs controls the detection of dietary protein, possibly by monitoring the entry of ingested amino acids into the TCA cycle. More broadly, these findings suggest a general strategy to screen for genes and pathways that might be used to boost the nutrient-regulated release of gut peptides such as GLP-1.

## Introduction

The small intestine is the principal site where ingested nutrients are directly sensed prior to absorption. This pre-absorptive nutrient sensing is mediated by enteroendocrine cells (EECs), a specialized class of sensory cells that are embedded within the intestinal epithelium and sample the chemical contents of intestinal lumen (Gribble and Reimann, 2019; Lu et al., 2021). Upon detection of nutrients or other chemical cues, EECs release into the circulation an array of hormones including CCK, GLP-1, GIP, PYY and 5-HT. These hormones can act locally on sensory neurons innervating the intestine or on distant organs to have a variety of effects on physiology and behavior, including changes in gastric emptying, intestinal motility, bile acid release, pancreatic function, and appetite. Importantly, modified forms of two of these hormones – GLP-1 and GIP – are now widely used as drugs to treat diabetes and obesity.

Nutrient detection by EECs triggers an increase in intracellular calcium that results in the exocytosis of hormone-containing vesicles. A number of molecules involved in this nutrient sensing process have been identified (Lu et al., 2021). For glucose and galactose, the most important sensor is thought to be Sodium-Glucose Transporter 1 (SGLT1, encoded by *Slc5a1*), a symporter that couples the transport of these sugars to sodium (Gorboulev et al., 2012; Gribble et al., 2003; Parker et al., 2012; Reimann et al., 2008). Thus, after a meal, elevated glucose and galactose in the small intestine drive sodium influx into EECs, resulting in EEC depolarization and calcium influx through voltage-gated calcium channels. It has also been proposed that intracellular glucose metabolism in EECs can be sensed independently through K-ATP potassium channels (Mace et al., 2012; Reimann et al., 2008; Reimann and Gribble, 2002), which may amplify the response to SGLT1-induced depolarization (Kuhre et al., 2015). For the detection of ingested fat, several molecules have been identified, including the fatty acid sensors *Gpr40* and *Gpr120* (Briscoe et al., 2003; Edfalk et al., 2008; Hirasawa et al., 2005) and the receptor *Gpr119*, which is activated by lipid metabolites such as oleoylethanolamide (Lauffer et al., 2009; Sihag and Jones, 2018).

In contrast to sugar and fat, the mechanisms by which EECs sense ingested protein are less well established (Hjørne et al., 2022). The two molecules that have been the most studied are *Slc15a1* (PEPT1), a symporter that couples the transport of di- and tri-peptides to protons (Diakogiannaki et al., 2013; Dranse et al., 2018), and the calcium-sensing receptor (*Casr*), a GPCR that is activated by the binding of Calcium as well as aromatic amino acids such as phenylalanine and tryptophan (Conigrave et al., 2007, 2000; Diakogiannaki et al., 2013; Modvig et al., 2019; Pais et al., 2016). There is evidence that these two proteins play a role in some responses to protein infusion into the small intestine (Alamshah et al., 2017; Dranse et al., 2018; Li et al., 2024). However, their restricted substrate specificity means that they cannot explain the ability of most individual amino acids to stimulate hormone release from EECs (Modvig et al., 2021). While several other sensors have been proposed, including *Gprc6a*, *Gpr142*, *Lpar5* (GPR93), *Slc38a2* (SNAT2) and the umami receptor *Tas1r1/r3*, there is conflicting evidence for and against the physiologic relevance of all these candidates (Choi et al., 2007; Clemmensen et al., 2017; Daly et al., 2013; Haid et al., 2011; Hjørne et al., 2022; Modvig et al., 2021; Oya et al., 2013; Rudenko et al., 2019; Wang et al., 2011; Wellendorph et al., 2005; Young et al., 2010).

All of these putative nutrient sensors in EECs were identified using candidate-based approaches, meaning they were tested based on their known biochemical function or known role in another system such as the pancreatic beta cell. To date, there have been no unbiased screens to identify genes expressed in EECs that are necessary for nutrient sensing and/or hormone release. We reasoned that such a screen might reveal not only novel candidate receptors, but also biochemical pathways and cellular processes that are important for nutrient sensing in general. We therefore set out to develop a strategy to screen for genes necessary for EEC nutrient sensing and applied this to investigate the detection of dietary protein.

## Results

To develop a strategy to screen for genes required for nutrient sensing in EECs, we took advantage of the fact that nutrient-detection in EECs triggers an increase in cytosolic calcium, which is the key step in driving hormone release. Although this nutrient-triggered calcium is transient, it can be converted into a stable fluorescent signal by using the reporter CaMPARI (Fosque et al., 2015), which undergoes a green-to-red transition in the presence of both calcium and UV light. Recently, it was shown, in neurons, that this ability of CaMPARI to irreversibly tag cells with elevated calcium can be used as the basis for a screen to identify genes required for calcium influx (Boggess et al., 2024). We therefore sought to adapt this screening approach for use in EECs (**Fig. 1a**).

**Figure 1.**
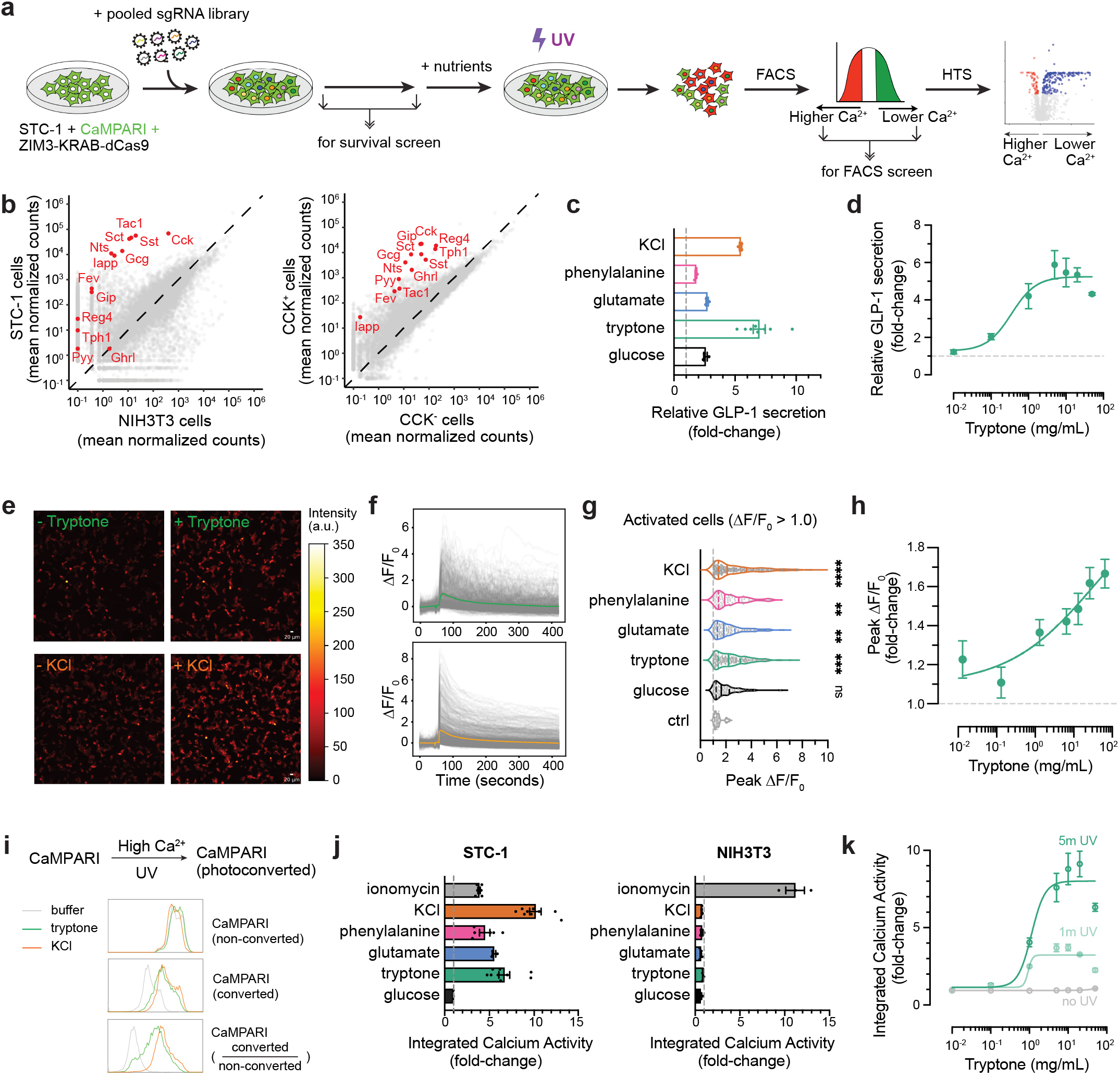
STC-1 cells secret GLP-1 and couple nutrient detection with calcium activity. a, Schematic of activity-based pooled CRISPRi screening. b, Gene expression levels for EEC and non-EEC cells determined by RNA-seq. EEC hallmark genes, including genes encoding secreted hormones, are highlighted in red. c-d, Relative GLP-1 secretion levels in STC-1 by ELISA. c, nutrient and non-nutrient (KCI) stimulation. d, Dose response curve for tryptone (amino acids and short peptides). e-h, Calcium activity by live cell GCaMP imaging. e, representative images of STC-1 stimulated by tryptone and KCI. f, .A.FIF_0_ traces for STC-1 cells after tryptone (top) or KCI (bottom) stimulation. g, Peak .A.FIF_0_ values for all activated cells. h, Dose response curve for tryptone. i-k, Integrated calcium activity by CaMPARI. i, flow cytometry analysis of CaMPARI photoconversion after different stimulation in STC-1. j, Median integrated calcium activity for STC-1 (left) and NIH3T3 (right) after nutrient and non-nutrient stimulation. ionomycin is a positive control that allows nonspecific calcium influx. k, Dose response curve for tryptone with various UV photoconvertion windows. *P < 0.05; **P < 0.01; ***P < 0.001; ****P < 0.0001.

Briefly, this strategy consists of: (1) Infecting EECs with a pooled library of thousands of sgRNAs, thereby enabling the knockdown of one gene at a time in each infected cell; (2) exposing this library of cells to a nutrient stimulus plus UV light, thereby tagging the cells that exhibit nutrient-induced calcium influx; (3) sorting these cells by flow cytometry to separate the nutrient-responsive from non-responsive cells; and (4) sequencing these two populations of cells to determine which sgRNAs are enriched in the cells that fail to respond to the nutrient stimulus. In the sections that follow, we describe the validation of each step in this approach.

### STC-1 cells detect nutrients and release GLP-1 *in vitro*

We first sought to identify an appropriate cell model for the screen. While primary EECs can be isolated and cultured, they are impractical for a high-throughput screen due to their low abundance (<1% of intestinal epithelium) and limited survival when cultured *in vitro* (<1 week). Aside from primary EECs, a number of immortalized EEC cell lines have been described, including STC-1, GLUTag, and Hu-80 cells. We tested these lines for nutrient-induced calcium influx and hormone release and found that STC-1 cells, a widely-used mouse cell line, exhibited the most robust responses.

Clonal populations of STC-1 cells can exhibit spontaneous heterogeneity in gene expression (Ryu et al., 2018). We therefore first characterized the STC-1 cells that we were using by RNA-Seq, which confirmed that they expressed hallmark EEC genes including nutrient-regulated peptides (*Cck*, *Gcg*, *Pyy*, *Sct*, *Nts*, *Tac1*, and *Gip*) as well as EEC markers like the transcription factor *Fev* (**Fig. 1b, left**), whereas most of these genes were absent in a control cell line (NIH-3T3 cells). Importantly, the same set of genes was enriched when we performed RNA-Seq of native *Cck*+ EECs isolated from the small intestine, and then compared this to *Cck*- cells from the same prep (**Fig. 1b, right**).

We next tested whether these STC-1 cells exhibit nutrient-triggered release of the hormone GLP-1 (**Fig. 1b-c**). Cells were incubated in a low nutrient buffer and then stimulated by addition of purified nutrients for 120 min, at which point the supernatant was removed and GLP-1 levels analyzed by ELISA (**Fig. 1c**). Compared to no stimulus control, GLP-1 release was induced by addition of glucose (1 mM - 2.6-fold increase) as well as tryptone (5 mg/mL – 7.0 fold increase), a protein hydrolysate that contains amino acids and short peptides and mimics dietary protein. The response to tryptone was dose-dependent with an EC_50_ of 0.35 mg/mL (**Fig. 1d**).

Consistent with this tryptone response, we also observed GLP-1 release in response to individual amino acids such as glutamate (2.7-fold) and phenylalanine (1.8 fold). These responses to nutrients could be mimicked by treatment with KCl, which directly depolarizes EECs and induced substantial GLP-1 secretion (5.5-fold; **Fig. 1c**). We also showed that the GLP-1 release in response to tryptone and KCl was unaffected by inclusion of glucose (1 mM) in the starvation buffer, indicating that complete nutrient deprivation is not required for these responses (**Fig. S1a**). Thus, STC-1 cells robustly release GLP-1 in response to nutrient stimulation as well as artificial depolarization.

We next sought to confirm that STC-1 cells exhibit increases in intracellular calcium in response to nutrient stimuli. We generated an STC-1 line that stably expresses GCaMP6m and then exposed these cells to the same nutrient panel used above and measured calcium-induced fluorescence by live imaging (**Fig. 1e-g**). Nutrient treatment caused a rapid increase in intracellular calcium which then slowly decayed over several minutes (**Fig. 1f**). This increase in calcium was significantly different from buffer treatment for all nutrients tested as well as KCl (**Fig. 1g**) and was dose-dependent for tryptone (**Fig. 1h**). This confirms that STC-1 cells can respond to nutrients with appropriate calcium influx.

The fact that STC-1 cells respond robustly to tryptone and amino acids suggests that they could be used to screen for mechanisms of protein sensing. To clarify whether the nutrient response in our STC-1 cells involves known mechanisms, we examined our RNA-Seq data to determine the expression of genes previously implicated in protein sensing. Surprisingly, we found that most of these putative protein sensors were not detectably expressed in our STC-1 line. This includes *Casr*, *Slc15a1* (i.e. PEPT1), *Slc15a2*, *Gprc6a*, *Gpr142*, *Lpar5* (i.e. GPR93), and *Tas1r1/3*. We independently confirmed the lack of expression of these genes by qPCR. The lone exception was *Slc38a2* (SNAT2), a sodium-coupled transporter for small neutral amino acids such as glutamine and alanine. However, when we knocked down *Slc38a2* expression by CRISPR interference (CRISPRi), there was no effect on the calcium response to tryptone or alanine and only a modest inhibition of responses to glutamine (**Fig. S1b**). These data imply that there exists an additional mechanism in STC-1 cells for sensing dietary protein and converting this into an electrical response and GLP-1 release. Importantly, this mechanism does not require, alone or in combination, any of the molecules that have previously been proposed to mediate protein sensing in EECs.

### CaMPARI can be used to permanently tag nutrient-responsive cells

CaMPARI undergoes photoconversion from green to red in the presence of both elevated calcium and UV light. This can be used to tag nutrient responsive cells as red and then separate them by FACS, a key step in our pooled screening approach (**Fig. 1a**). We therefore optimized CaMPARI photoconversion for detecting nutrient-induced calcium activity in EECs.

We first generated an STC-1 line that stably expresses CaMPARI and measured by flow cytometry the efficiency of CaMPARI photoconversion after nutrient or KCl stimulation. We found that treatment with tryptone or KCl, followed by brief UV light treatment, resulted in a clear increase in the percentage of photoconverted (red) cells (**Fig. 1i, Fig. S1d**). Calcium was both necessary and sufficient for this increase in photoconversion, as the calcium ionophore ionomycin could substitute for nutrient stimulation (**Fig. 1j**) whereas the response to nutrients was completely blocked by the inclusion of the calcium chelator EDTA (**Fig. S1e**).

Photoconversion was also dependent on the nutrient-sensing machinery within STC-1 cells, as we saw no photoconversion upon treatment of a control cell line (NIH-3T3 cells) with either nutrients or KCl (**Fig. 1j, left**), although these cells could respond when ionomycin was used to drive non-specific calcium entry (**Fig. 1j, right**). The photoconversion also depended on UV light exposure, which was dose-dependent (**Fig. 1k**) with effective labelling observed after 5 min.

Thus, CaMPARI can be used to faithfully tag STC-1 cells based on their calcium response to nutrients. In the remainder of the manuscript, we use the extent of CaMPARI photoconversion (i.e. the ratio of converted (red) to non-converted (green) cells, which we refer to as integrated calcium activity) as a readout for this calcium response.

Because we are interested in identifying mechanisms for sensing dietary protein, we next characterized the ability of CaMPARI to read out responses to protein specifically. First, we noted that the dose-response profile for tryptone-induced CaMPARI photoconversion (**Fig. 1k**, EC_50_: 0.93 mg/mL at 1 min UV and 1.13 mg/mL at 5 min UV) was similar to the dose-response for tryptone-induced GLP-1 secretion (**Fig. 1d**, EC_50_: 0.35 mg/mL), consistent with the idea that CaMPARI is measuring intracellular calcium which stimulates GLP-1 release. Next, we tested the ability of individual amino acids to induce CaMPARI photoconversion in STC-1 cells. We found that treatment with most amino acids induced a measurable increase in photoconversion (**Fig. S1c**), with small, uncharged amino acids generally yielding the strongest responses (e.g. Ser, Pro, Gly, Val, Thr, Ala, Asn, Gln) and large or charged amino acids generally yielding weaker responses (e.g. Tyr, Arg, Ile, Asp, Trp, Lys). Thus, CaMPARI can capture STC-1 responses to either a protein mixture (tryptone) or individual amino acids.

### Pilot screens identify specific modulators of amino acid sensing in STC-1 cells

To enable the use of CRISPRi to screen for modulators of amino acid sensing, we next made an STC-1 cell line that stably expresses both CaMPARI and ZIM3-KRAB-dCas9, which is a potent transcriptional repressor fused to dCas9 to enable CRISPRi knockdown of gene expression (Alerasool et al., 2020). We tested multiple clonal lines and chose clone 2C6 due to the fact that it exhibited high and consistent CRISPRi KD efficiency (**Fig. S2a-b**).

To perform a pilot screen as outlined in **Fig. 1a**, we first obtained a medium-scale mouse sgRNA library (CRISPRi_v2 sublibrary m1; Addgene #83989). This library contains 12,100 sgRNAs targeting 2,269 genes (primarily kinases, phosphatases and drug targets), as well as 250 non-targeting control sgRNAs. We transduced the STC-1 cell clone 2C6 with this lentiviral sgRNA library at a low MOI to ensure no more than one sgRNA was introduced in any cell. We then selected for positively transduced cells with puromycin and also confirmed infection by measuring BFP fluorescence. After allowing at least six cell doublings, cells were treated with the stimulus (nutrient or control), followed by UV light for 5 min to convert the transient calcium response to permanent red fluorescence. Cells were then immediately harvested and sorted by FACS to separate the responsive (high red fluorescence, low green fluorescence) from non-responsive (high green, low red) cells (**Fig. S2c**). Sequencing of genomic DNA from the sorted cells allowed us to quantify the sgRNAs that were enriched in each population.

We performed this screen in response to two amino acid stimuli (tryptone and phenylalanine) and, as a control, one non-nutritive stimulus that induces robust calcium influx (KCl). In addition, we separately analyzed the sgRNA diversity in cell samples collected on day 0 and the day before FACS sorting. This “dropout screen” served as an in-screen control for changes in cell survival induced by specific sgRNAs.

The results of these screens are depicted in **Fig. 2** as volcano plots (**Fig. 2a-c**, left). We also calculated a metric, which we called phenotype score, which is defined as the product of the effect size and significance (i.e. the product of the two axes in the volcano plot). We then ranked genes by their phenotype scores and labeled the top 10 hits from each screen (**Fig. 2a-c**, right). Overall, approximately 4-5% of targeted genes scored as hits (FDR < 0.05) in the tryptone and phenylalanine screens (tryptone: 93 positive hits, 12 negative hits; phenylalanine: 81 positive hits, 13 negative hits). The fact that most hits were positive regulators of EEC activation (i.e. genes that, when knocked down, decrease nutrient-induced calcium, labelled in blue) rather than negative regulators (i.e. genes that, when knocked down, increase nutrient-induced calcium, labelled in red) may be due to the fact that we screened with high concentrations of tryptone, creating a ceiling effect.

**Figure 2.**
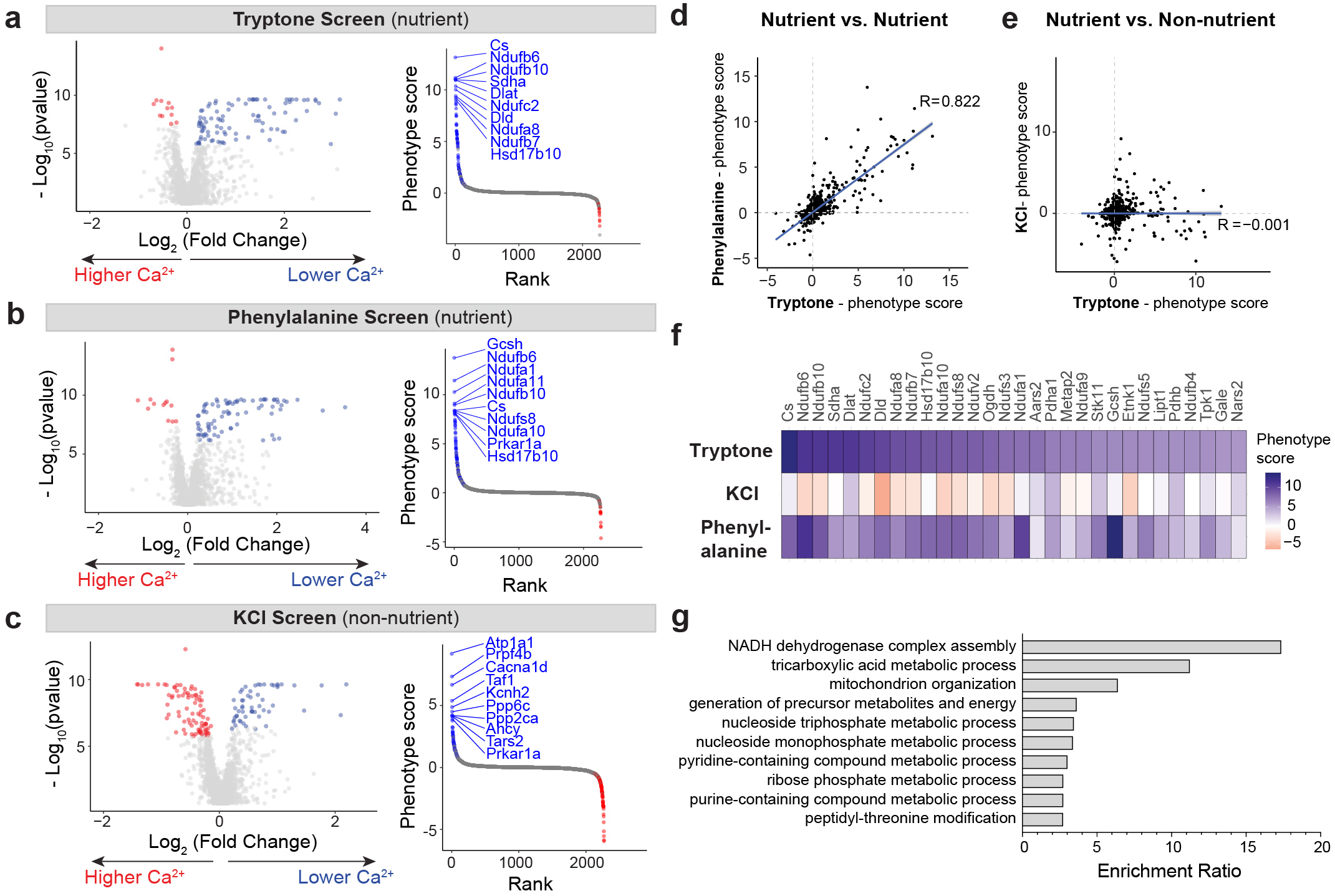
Parallel pooled CRISPRi screens identify modulators specific to amino acid sensing in STC-1 cells. a-c, Volcano plots and rank plots for parallel screens performed with Weissman’s m1 subpooled library.band care nutrient screens, whiled is a non-nutrient screen. Significant hits (FDR< 0.05) are highlighted in volcano plot, and top 10 hits with highest phenotype scrore [log_2_(fold change) x -log_10_(pvalue)] are highlighted in rank plots. d-e, Phenotype scores of all library genes from two parallel screens. d, Tryptone (nutrient) vs. phenylalanine (nutrient). e, Tryptone (nutrient) vs. KCI (non-nutrient). f, Heatmap showing top 30 hits from the tryptone screen. g, Pathway enrichment analysis of all positive hits.

We reasoned that if the screens worked as intended, then the hits identified in the tryptone and phenylalanine screens should be similar to each other, because both screens involve protein-induced EEC activation. On the other hand, the hits identified in the KCl screen should be different, because KCl and protein utilize different mechanisms to cause EEC activation. To test this, we compared the phenotype scores for all targeted genes between these pairwise combinations of screens. Strikingly, this revealed that the phenotype scores from the nutrient screens were highly correlated (**Fig. 2d**; Pearson correlation coefficient R = 0.822), whereas there was no correlation between the tryptone screen and the KCl screen (**Fig. 2e**; Pearson correlation coefficient R = -0.001). This overlap was particularly clear when we examined the top 30 hits from the tryptone screen: virtually all of these genes were top hits from the phenylalanine screen, and none were hits in the KCl screen (**Fig. 2f**). We observed the same trend when we examined the enrichment at the level of individual sgRNAs rather than genes (**Fig. S2d**).

Importantly, there was no correlation when we compared the hits from the protein screens with the hits from the internal survival screen, indicating that we did not simply select for sgRNAs that are toxic (**Fig. S2e-g**; tryptone screen, R = -0.26; phenylalanine screen, R = -0.27). Taken together, these data indicate that the hits identified in these screens are linked to stimulus-specific mechanisms of EEC activation.

To understand the function of the genes that were enriched in the protein screens, we performed pathway enrichment analysis for all the positive regulators. This revealed that mitochondrial energy metabolism pathways were overwhelmingly enriched among the genes required for protein sensing (**Fig. 2g**). This includes components of the electron transport chain and genes involved in the tricarboxylic acid (TCA) cycle. For example, we identified numerous components of the NADH dehydrogenase complex (Complex I) as hits in the tryptone and phenylalanine screens. Given that amino acids enter the TCA cycle and are metabolized within mitochondria, this suggests a possible mechanism for linking intracellular amino acid metabolism to protein sensing.

### A large-scale screen identifies mitochondrial metabolic pathways as essential for amino acid sensing

Encouraged by these results, we designed a larger screen with a custom library targeting 3,744 mouse genes (**Fig. 3a**). This includes most receptors (GO: 0004888), transmembrane transporters (GO: 0022857) and metabolic enzymes (GO: 0006520, 0022900, 0046031, 0006091), the three functional categories that we reasoned are most likely to contain modulators of amino acid sensing. We also included all of the hits from the pilot screens and the top 150 differentially enriched genes in STC-1 cells compared to NIH3T3 cells from our RNA-seq data, while we removed any genes not expressed in our STC-1 cells. Each gene was targeted by the top five sgRNAs from the mouse CRISPRi v2 library, and we also included 490 non-targeting control sgRNAs (∼2.5%), resulting in a total of 20,250 sgRNAs (**Fig. 3a**). Of note, the scale of this screen was determined by the fact that, in practice, ∼ 20,000 elements is the upper limit if we wish to complete cell sorting within a single day and achieve > 500x coverage for each FACS-sorted population.

**Figure 3.**
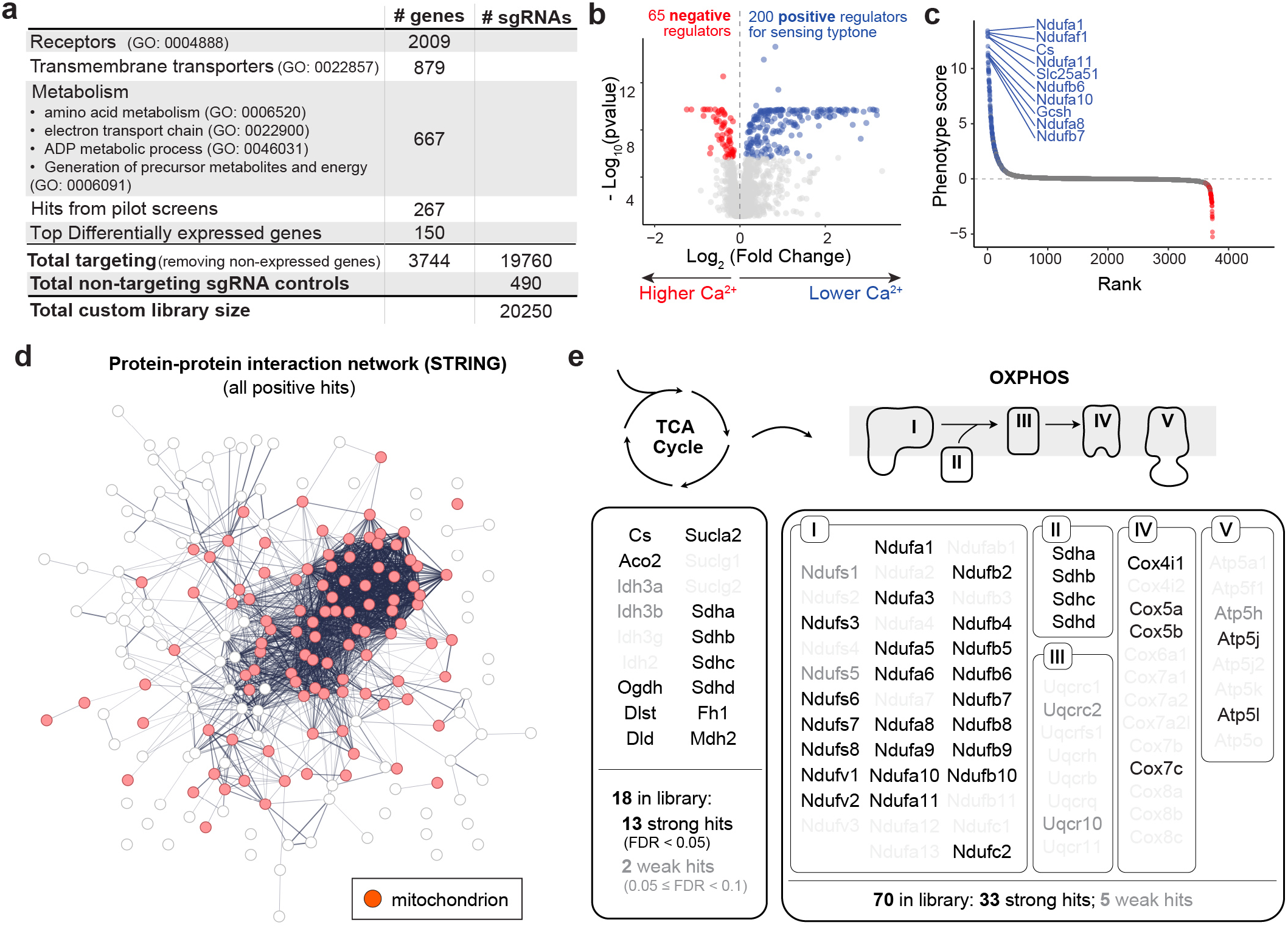
Large-scale screen with custom library identifies mitochondrial energy metabolism as essential for amino acid sensing. a, Composition of custom sgRNA library for large-scale CRISPRi screening. b-c, Volcano plot and rank plot for custom library screen. b, Significant hits (FDR < 0.05) are highlighted. c, Top 10 hits with highest phenotype scrore [log_2_/fold change) x -log_10_ (pvalue)] are highlighted. d, Functional protein-protein interaction network for all positive hits by STRING. Line thickness indicates the strength of data support for interaction. Genes with mitochondrial annotation (GO:0005739) are highlighted in red. e, Hit distribution for two most critical mitochondrial energy metabolism pathways, TCA cycle and OXPHOS. Strong hits with FDR< 0.05 are highlighted in black, and weak hits with FDR< 0.1 are labeled in ‘gray50’. Non-hit genes with FDR ≥ 0.1 are ‘gray10’.

We used this library to perform a screen with tryptone as the stimulus, which identified 200 genes as positive regulator hits (i.e. genes for which knockdown decreased nutrient-induced calcium) and 65 as negative regulators. This data is presented as volcano plot (**Fig. 3b**) and as a list ranked by phenotype score with the Top 10 hits highlighted (**Fig. 3c**). We noted, first, that there was a high degree of reproducibility between the pilot and large-scale screens. That is, for the subset of genes targeted in both screens, there was a high correlation between their phenotype scores in the two screens, both for tryptone-induced calcium activity (**Fig. S3b**, Pearson correlation coefficient R = 0.87) and for survival (**Fig. S3a**, R = 0.96). Moreover, the efficiency of CRISPRi knockdown in the large scale screen was validated by the high recall rate (0.793 at phenotype score threshold of 1) for common essential genes evaluated by the survival screen (**Fig. S3d**). However, there was again no correlation between the phenotype score in this survival screen and the phenotype score in response to tryptone (R=-0.39), indicating that we are not selecting for toxic genes (**Fig. S3**).

Consistent with the pilot screens, pathway enrichment analysis revealed that the positive hits are highly enriched in mitochondrial metabolic pathways (**Fig. S3e**). Indeed, over half of the positive hits (103/200 genes) are expressed in the mitochondria and directly involved in energy metabolism. We also found, through functional protein-protein interaction network analysis by STRING, that the top hits are derived from a core interaction group composed of mostly TCA cycle and OXPHOS genes (**Fig. 3d** and **Fig. S3f**). For example, out of the 18 genes involved in the TCA cycle that were included in our custom library, 13 emerged as strong hits (FDR < 0.05), and 2 as weak hits (FDR < 0.1) (**Fig. 3e**). For OXPHOS, out of 70 genes included in our library, 33 emerged as strong hits and 5 as weak hits. This included genes involved in all five complexes of the electron transport chain, including most prominently in Complex I and II (**Fig. 3e**). In addition to genes involved in respiration, top hits included a number of enzymes involved in glycolysis (*Gpi1*, *Aldoa*, *Tpi1*, *Pgk1*) as well as a number of mitochondrial aminoacyl-tRNA synthetases (*Cars2*, *Pars2*, *Sars2*, *Ears2*), among other gene families.

### Mitochondrial respiration bidirectionally modulates nutrient sensing in EECs

We next sought to validate individual hits from the screen, focusing on the genes related to mitochondrial respiration. We selected two genes from the top 10 hits in the screen, *Cs* (citrate synthase; TCA cycle) and *Ndufb6* (a component of OXPHOS Complex I) (**Fig. 4a**), and knocked them down individually in STC-1 cells and evaluated calcium responses and GLP-1 secretion. We first validated the efficiency of knockdown (**Fig. S2b**, left; *Cs*, 93.1 ± 2.8% KD; *Ndufb6*, 95.3 ± 3.9% KD) and further showed that this CRISPR-induced depletion of *Cs* or *Ndufb6* had no effect on baseline calcium activity (*Cs*, 0.943 ± 0.098, p = 0.8174; *Ndufb6*, 0.900 ± 0.112, p = 0.6464, 2-way ANOVA with Dunnett’s multiple comparisons test). However, knockdown of these genes completely blocked the tryptone-induced calcium response (**Fig. 4b**; NTC: 9.128 ± 0.841; *Cs*: 1.093 ± 0.071, p = 0.0038; *Ndufb6*: 0.993 ± 0.064, p = 0.0037). We observed a similar ability of gene knockdown to block tryptone-induced calcium in validation experiments with eight other top hits (**Fig. S4a**). Consistent with the reduction in calcium, loss of *Cs* or *Ndufb6* also completely abolished tryptone-induced GLP-1 secretion (**Fig. 4c**; NTC: 5.383 ± 0.684; *Cs*: 0.567 ± 0.032, p < 0.0001; *Ndufb6*: 1.031 ± 0.091, p < 0.0001, 2-way ANOVA with Dunnett’s multiple comparisons test). This confirms that the TCA cycle and OXPHOS play an essential role in amino acid sensing in EECs.

**Figure 4.**
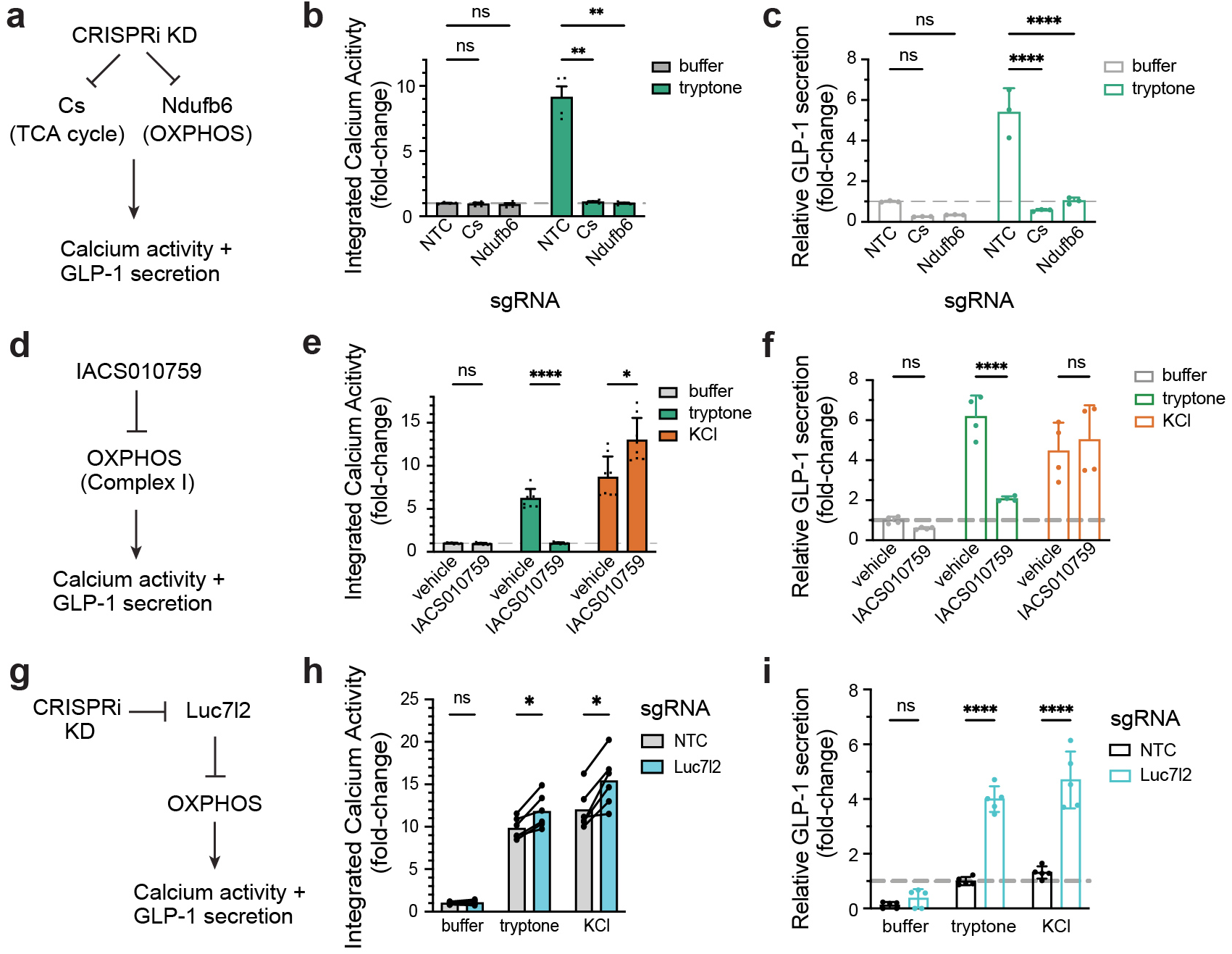
Mitochondria respiration bidirectionally modulates nutrient sensing. a-c, Valiation of top hits in mitochondrial respiration pathways by CRISPRi KD. a, Schematic for experimental design. b, Integrated calcium activity in STC-1 stably expressing non-targeting control (NTC) or sgRNA targeting top hit genes. c, Relative GLP-1 secretion in STC-1 after CRISPRi KD. d-f, Validation of the role of mitochondrial respiration in amino acid sensing by pharmacological inhibition of OXPHOS Complex I. d, Schematic for experimental design. e, Integrated calcium activity in STC-1 cells pretreated with vehicle or IACS010759. f, Relative GLP-1 secretion in STC-1 after stimulation, with vehicle or IACS010759. g-i, Stimulating OXPHOS boosts EEC activity and GLP-1 secretion. g, Schematic for experimental design. h, Integrated calcium activity in STC-1 stably expressing NTC or sgRNA targeting Luc712, an inhibitor of OXPHOS. i, Relative GLP-1 secretion in STC-1 with the indicated perturbation and stimulation. *P < 0.05; **P < 0.01; ***P < 0.001; ****P < 0.0001.

The experiments above utilize a genetic perturbation that causes a chronic reduction in mitochondrial respiration, which could have many effects on cell physiology. To test the acute requirement for mitochondrial respiration in EEC protein sensing, we used a pharmacological inhibitor (IACS010759, a potent and selective inhibitor of OXPHOS Complex I) to block respiration immediately before nutrient stimulation. We first performed a dose-response, which demonstrated that IACS010759 treatment results in dose-dependent inhibition of tryptone-induced calcium activity (**Fig. S4b**) with nearly complete inhibition at 100 nM IACS010759 (**Fig. 4e**; vehicle 6.190 ± 0.395 vs. IACS010759 0.999 ± 0.035, p < 0.0001; 2-way ANOVA with Šidák’s multiple comparisons test). We then performed a timecourse in which we preincubated IACS010759 with STC-1 cells for various durations (0-60 min) prior to nutrient stimulation and then measured tryptone-induced photoconversion. We found that IACS010759 already reduced the nutrient response at the earliest timepoint we could measure (0 min, i.e. adding the drug and nutrient simultaneously; **Fig. S4c**) and that this inhibitory effect strengthened within the first few minutes of incubation (e.g. 19.6% of control with 1 min incubation, 11.6% with 5 min, and 8.5% with 10 min). Moreover, we confirmed using a Seahorse Analyzer that IACS010759 can inhibit OXPHOS in STC-1 cells on this timescale (**Fig. S4d**). Thus, mitochondrial respiration is acutely required for EEC activation by amino acids, which suggests that respiration may be part of the proximal sensing mechanism.

We reasoned that if mitochondrial respiration is a specific mechanism for sensing amino acids, then blocking OXPHOS should not inhibit the calcium response to KCl. Indeed, we found that IACS010759 treatment *increased* the calcium activity in response to KCl (**Fig. 4e**; vehicle 8.670 ± 0.856 vs. IACS010759 12.963 ± 0.917, p = 0.0124). Of note, this argues that STC-1 cells are not generally functionally impaired by IACS010759 treatment on this timescale. We also tested the effect of IACS010759 on release of GLP-1 and found that, similar to the results with calcium, IACS010759 blocked tryptone-induced GLP-1 secretion (**Fig. 4f**, vehicle 6.176 ± 0.527 vs. 2.068 ± 0.060, p < 0.0001), but did not prevent GLP-1 release in response to KCl (**Fig. 4f**, vehicle 4.453 ± 0.710 vs. 5.013 ± 0.862, p = 0.4445, 2-way ANOVA with Tukey’s multiple comparisons test). Taken together, these data show that mitochondrial respiration is acutely required for tryptone-induced calcium influx and GLP-1 release but is dispensable for these same effects after KCl stimulation. This strongly argues there is a specific mechanistic link between respiration and amino acid detection.

If this connection between respiration and amino acid detection is direct, then we would predict that it should also be bidirectional: that is, increasing respiration should increase tryptone-induced calcium activity and GLP-1 secretion. While there is a paucity of good methods for boosting the rate of respiration in vitro, a screen for genes that control mitochondrial function identified the gene Luc7l2 as an inhibitor of OXPHOS (Jourdain et al., 2021). We therefore knocked this gene down in STC-1 cells by CRISPRi and assessed the effect on nutrient-induced responses (**Fig. 4g**). Consistent with our hypothesis, we found that when Luc7l2 was knocked down there was an increase in both calcium activity (**Fig. 4h**; tryptone, q = 0.0032; KCl, q = 0.0032; multiple paired t tests) and GLP-1 secretion (**Fig. 4i**; tryptone, p < 0.0001; KCl, p < 0.0001; 2-way ANOVA with Šidák’s multiple comparisons test) in response to both tryptone and KCl. This indicates that mitochondrial respiration is a bidirectional regulator of the pathways that lead to calcium influx and hormone release from EECs, with gain-of-function manipulations broadly potentiating these responses and loss-of-function specifically preventing nutrient-induced activation.

### Amino acid sensing in EECs involves the entry of amino acids into TCA cycle

We next set out to examine the mechanisms underlying the connection between mitochondrial respiration and amino acid detection. Given that amino sensing by EECs requires genes involved in the TCA cycle and OXPHOS (**Figs. 2** and **3**), and given that this requirement is acute and specific to nutrients (**Fig. 4**), we reasoned that amino acids may be detected via a byproduct of their metabolism and subsequent entry into TCA cycle.

To test this, we sought to identify a key enzyme that is a choke point for the entry of a specific amino acid into the TCA cycle and then examine the effect on nutrient-induced calcium activity when this enzyme is disrupted. We found that glutamine induced robust calcium activity in EECs (**Fig. S1c**) and the breakdown of glutamine into glutamate is an essential step for this nutrient to enter the TCA cycle (**Fig. 5a**). The metabolism of glutamine into glutamate is performed by two enzymes, *Gls* and *Gls2*, the latter of which is not expressed in STC-1 cells. Thus, knockdown of *Gls* alone should be sufficient to prevent this conversion and glutamine metabolism. We therefore knocked down *Gls* by CRISPRi and, consistent with our hypothesis, we found that this greatly reduced the calcium response to glutamine treatment (**Fig. 5b**; glutamine, p < 0.0001, 2-way ANOVA with Šidák’s multiple comparisons test). As controls, we tested amino acids that are metabolized by different routes (proline and glutamate) as well as tryptone, which contains all 20 amino acids. For none of these control stimuli did *Gls* knockdown affect the calcium response (**Fig. 5b**; proline, p = 0.4734; glutamate, p = 0.8287; tryptone, p = 0.8391; 2-way ANOVA with Šidák’s multiple comparisons test). Thus, the effects of glutamine on EEC calcium activity is blocked by preventing it from entering the TCA cycle.

**Figure 5.**
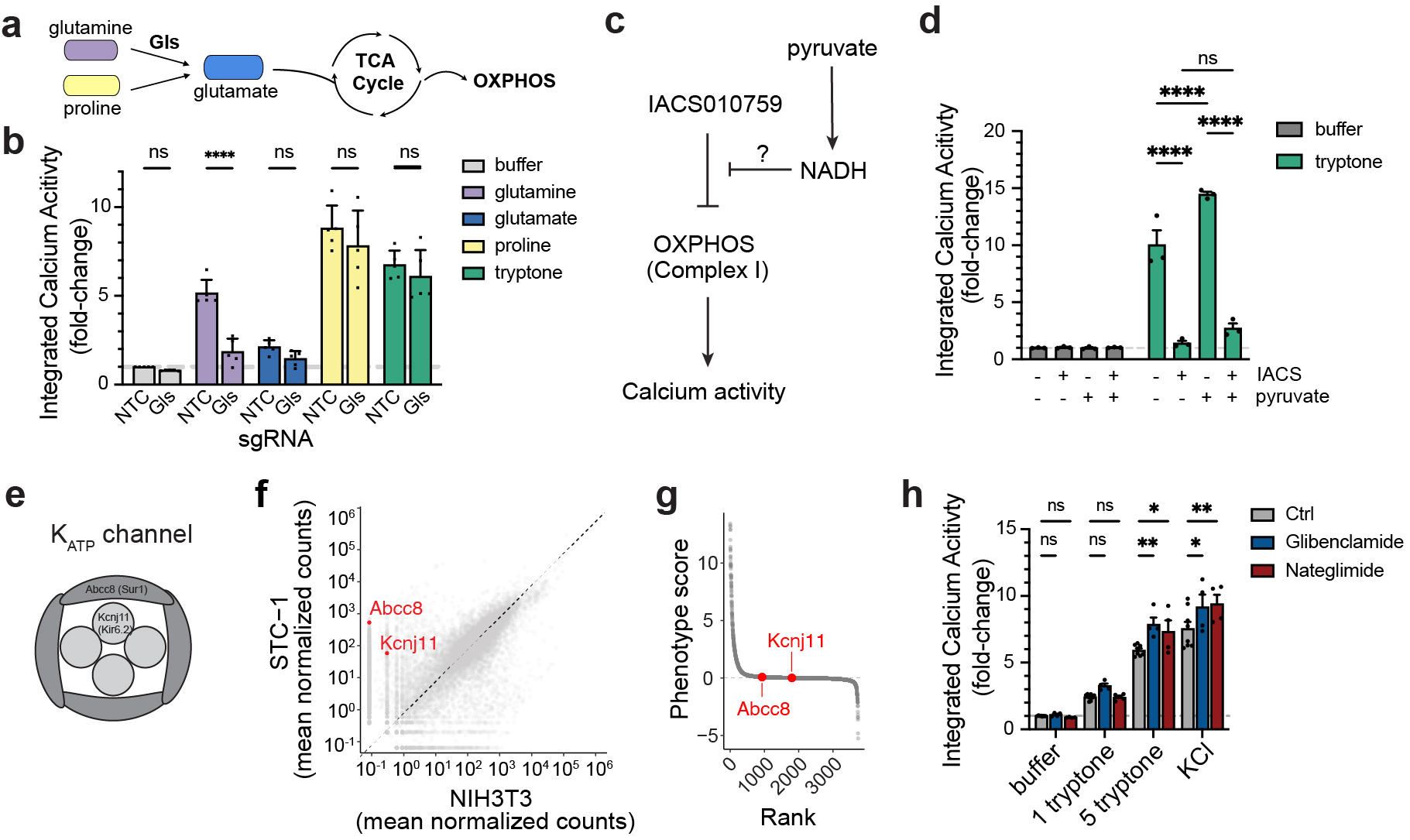
Amino acid sensing in EECs requires the entry into TCA cycle. a-b, Amino acid metabolism and entry into the TCA cycle is required for EEC sensing. a, Gls is a key enzyme required for glutamine metabolism and its entry into TCA cycle, but not for praline or glutamate. b, Integrated calcium activity in STC-1 stably expressing NTC or sgRNA targeting Gls. c-d, Restoring NADH and redox is not sufficient for amino acid sensing when OXPHOS is inhibited. c, Schematic for experimental design. d, Integrated calcium activity in STC-1 with the indicated treatments and stimulation. Cells were pre-treated with vehicle/lACS and/or pyruvate for 1 h before stimulation. e-h, K_ATP_ channel is dispensable for amino acid sensing in STC-1. e, Schematic of K_ATP_ channel, composed of Kcnj11 and Abcc8. f, Gene expression levels of Abcc8 and Kcnj11 in STC-1 vs. NIH3T3 by RNA-seq. g, Rank plot showed neither Abcc8 nor Kcnj11 is a hit from the custom library screen by tryoptone. h, pharmacological inhibition of K_ATP_ channel in STC-1 does not increase baseline calcium activity, but moderatly increase acitivty with strong stimulation (5 mg/ml tryptone or KCI). *P < 0.05; **P < 0.01; ***P < 0.001; ****P < 0.0001.

Manipulations of OXPHOS could affect cell physiology, and thereby signal nutrient detection, by causing changes in redox balance or changes in energy production. We first tested the role of redox balance. In this regard, inhibition of Complex I by the hits in our screen or IACS010759 would be predicted to prevent the oxidation of NADH to NAD+, leading to accumulation of NADH and an increase in reactive oxygen species. This can be counteracted by treatment with pyruvate, which drives cytosolic reactions that consume NADH and thereby restore the NAD+/NADH balance. We therefore tested whether pretreatment with pyruvate could rescue the inhibition of tryptone-induced calcium activity by IACS010759. However, we found that pyruvate treatment had no effect (**Fig. 5d**; p = 0.2761; 2-way ANOVA with Tukey’s multiple comparisons test). To test this a different way, we used heterologous expression of Ndi1, the yeast NADH dehydrogenase. This enzyme can convert NADH into NAD+ but does not drive ATP synthesis through proton pumping and therefore can be used to counteract redox imbalances (Seo et al., 1998). However, expression of Ndi1 did not significantly rescue the inhibition of tryptone-induced calcium activity by IACS010759 (**Fig. S4e**). Together, these data suggest that maintenance of redox balance is not the primary reason that OXPHOS is required for amino acid sensing.

An alternative possibility is that disruptions to OXPHOS cause changes in energy production, possibly by preventing the metabolism of amino acids, and that this results in changes in calcium influx and hormone release. The most well characterized mechanism for linking intracellular energy metabolism to hormone secretion is ATP-sensitive potassium (K_ATP_) channels, which are inhibited when ATP levels are high, resulting in cellular depolarization. This mechanism has been extensively studied in the context of the pancreatic beta cell and, to a lesser extent, in EECs (Kuhre et al., 2015; Reimann et al., 2008; Reimann and Gribble, 2002).

To explore whether K_ATP_ channels are responsible for amino acid sensing in STC-1 cells, we first confirmed that both components, *Abcc8* (SUR1) and *Kcnj11* (KIR6.2) (**Fig. 5e**), were expressed in STC-1 cells (**Fig. 5f**). However, neither gene was a hit from our tryptone screen, suggesting that they are not required for the EEC response (**Fig. 5g**). To confirm this screening result, we tested whether addition of two K_ATP_ inhibitors (Glibenclamide or Nateglimide) would modulate calcium influx. These drugs would be predicted to mimic a cellular state of nutrient and energy surplus, but they failed to induce calcium activity in STC-1 cells at baseline (**Fig. 5h**; buffer: Glibenclamide 1.093 vs. 1.000, p = 0.9825, Nateglimide 0.890 vs. 1.000, p=0.9754; 1 mg/mL tryptone: Glibenclamide 3.260 vs. 2.415, p = 0.2596; Nateglimide 2.400 vs. 2.415, p=0.9995). However, treatment with these drugs did produce a subtle increase in the calcium response following strong stimulation produced by either high concentrations of tryptone or KCl (**Fig. 5h** 5 mg/mL tryptone: Glibenclamide 7.858 vs. 5.920, p = 0.0027; Nateglimide 7.335 vs. 5.920, p=0.0322; KCl: Glibenclamide 9.165 vs. 7.544, p = 0.0127; Nateglimide 9.395 vs. 7.544, p=0.0042; 2-way ANOVA with Dunnett’s multiple comparisons test). This suggests that K_ATP_ channels do not play a central role in amino acid sensing in STC-1 cells, but they may tune the excitability of the cells to boost activation under strong stimulation.

### Mitochondrial respiration is essential for amino acid sensing in primary EECs

We next sought to validate that these effects we observe in STC-1 cells are also observed in primary EECs. To do this, we targeted expression of GCaMP8s to EECs by crossing CCK^Cre/+^ mice to mice expressing a Cre-dependent GCaMP8s reporter (Igs7^GCamp8s/+^). We then isolated intestinal epithelial cells from the small intestine of these mice and cultured them in a monolayer (**Fig. 6a**). After 3 days in culture, we performed live cell calcium imaging using tryptone and KCl as stimuli **(Fig. 6b**). To enable within-cell comparisons, cells were treated first with vehicle and sequentially exposed to tryptone and KCl, and then treated with IACS010759 and exposed to tryptone and KCl again. We found that, while vehicle (DMSO) caused a slight decrease in the calcium response to tryptone (tryptone normalized to KCl, DMSO p = 0.0104, paired t-test), IACS010759 completely inhibited the tryptone-induced calcium response without any effect on the KCl response (tryptone normalized to KCl, IACS010759 p < 0.0001; tryptone IACS010759 p = 0.0089; KCl IACS010759 p = 0.1340; paired t-test) (**Fig. 6b-e**). This effect was also observed when we analyzed the response to tryptone directly, without normalizing to the KCl response (**Fig. S5**). This confirms that mitochondrial respiration is acutely required for the activation of primary EECs by amino acids.

**Figure 6.**
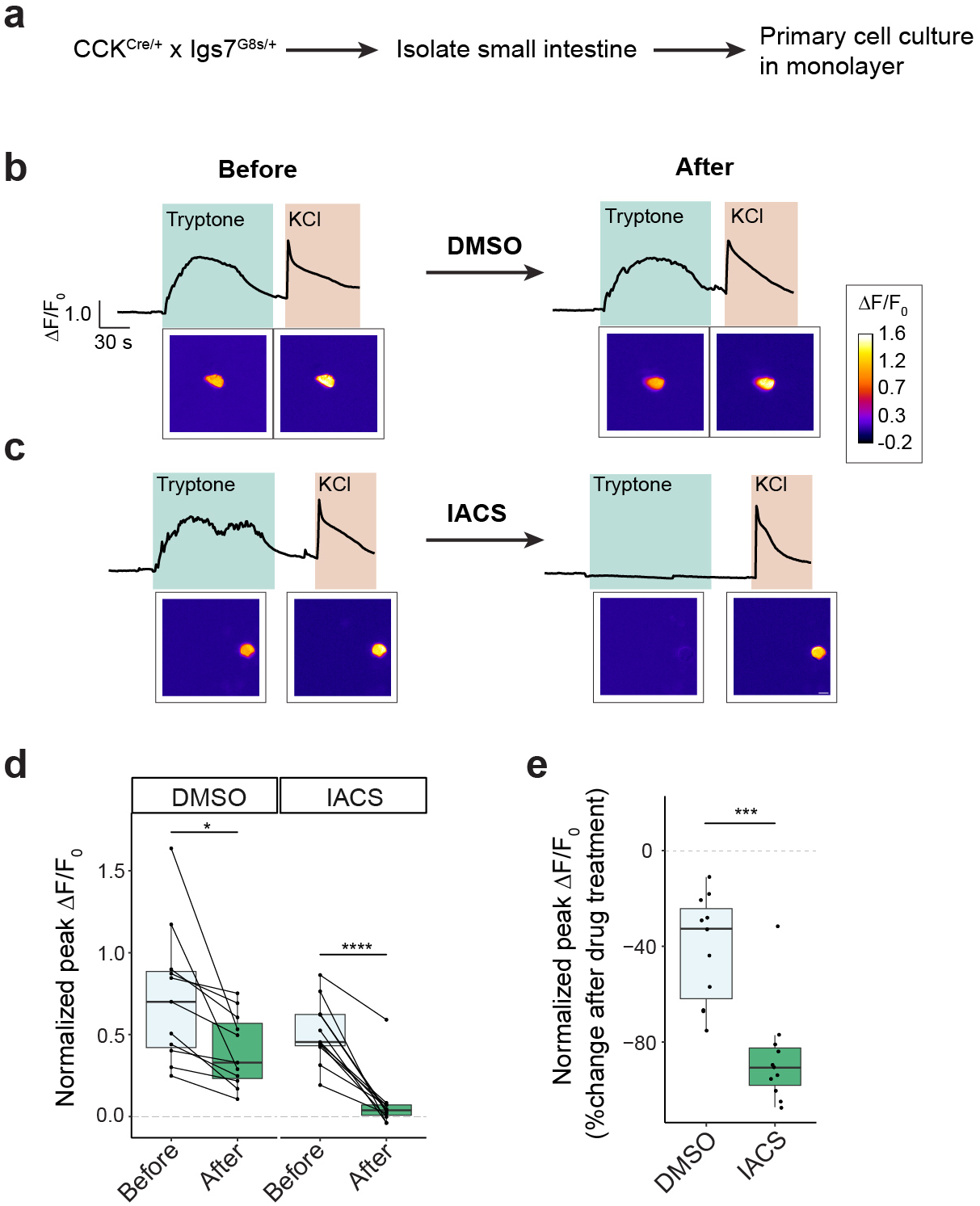
Mitochondrial respiration is essential for amino acid sensing in primary EECs. a, Schematic of experimental design. b-c, Representative GCaMP imaging ΔF/F_0_ traces and images comparing the same cell before and after treatment. b, DMSO control. c, IACS010759 (inhibitor of OXPHOS Complex I). d-e, Quantification of calcium activity in primary EECs before and after treatment. Pairwise comparison of same cells is shown. d, Peak ΔF/F_0_ for tryptone normalized by KCI. e, Percent of change after treatment for peak ΔF/F_0_for tryptone normalized by KCI. *P < 0.05; ***P < 0.001; ****P < 0.0001.

## Discussion

The molecular mechanisms for sensing of dietary protein in the intestine remain poorly understood. A particular challenge for protein sensing, relative to other macronutrients, is the chemical diversity of amino acids and short peptides that are known to be detected by the body and transduced into a hormonal response. While a number of putative protein and amino acid sensors have been described, these molecules alone are unlikely to fully explain the ability to detect ingested protein in the gut. Indeed, we found that both tryptone and individual amino acids elicited robust calcium responses and GLP-1 release in an EEC cell line that lacks expression of almost every known protein sensor. This indicates that another mechanism for protein sensing must exist.

To identify this mechanism, we set out to develop an unbiased approach to screen for genes that are necessary for nutrient sensing in EECs. Our screen takes advantage of the fact that nutrient stimulation causes depolarization and calcium influx, properties that have been used, in other contexts, as the basis for screens to identify receptors for other important physiologic processes (Caterina et al., 1997; Coste et al., 2010). While our initial expectation was that this screen would reveal novel transmembrane receptors that detect amino acids, we found instead that the overwhelming majority of hits in our screen were genes involved in OXPHOS and the TCA cycle. We confirmed in secondary assays that mitochondrial respiration is required for the response to protein, but not a non-nutritive stimulus, in both an EEC cell line and primary EECs. We further showed that this link between respiration and protein sensing is acute and bidirectional, and that preventing the entry of one amino acid – glutamine – into the TCA cycle is sufficient to prevent EEC activation by that nutrient. These data strongly suggest that EECs detect ingested amino acids, at least in part, by monitoring a byproduct of their metabolism in the mitochondria. Of note, this mechanism would provide a simple solution for how the cell is able to monitor the abundance of 20 chemically diverse amino acids, since all amino acids can be converted into intermediates that enter the TCA cycle.

Previous work has shown that mitochondrial respiration is important for gut metabolism and function. For example, mutations that impair mitochondrial function prevent enterocytes from properly processing ingested lipids into chylomicrons (Moschandrea et al., 2024). Although we are unaware of any studies that have directly measured how changes in mitochondrial function affect EEC hormone release in vivo, one study described the changes in gene expression that occur in the intestine in response to vertical sleeve gastrectomy (VSG) (Koch-Laskowski et al., 2024), a procedure that causes a dramatic increase in the release of hormones such as GLP-1 (Peterli et al., 2012). This study found that VSG was prominently associated with increased expression of genes involved in mitochondrial respiration. Thus, this provides correlative evidence that increased respiration in the intestinal epithelium is association with increased GLP-1 levels in vivo.

The development of drugs based on the incretins GLP-1 and GIP has revolutionized the treatment of diabetes and obesity. While these drugs are highly effective, they require administration of pharmacologic doses that increase the hormone concentration in the blood by >1,000-fold, and this is associated with significant GI side effects in many individuals. An alternative strategy for leveraging incretin biology would be to boost the natural release of these hormones from their endogenous sites in the intestine, which might have beneficial physiologic effects at much lower concentrations. Aside from DPP-4 inhibitors, which delay GLP-1 breakdown but do not boost its release, we are unaware of any targeted agents that stimulate hormone production from EECs as their mechanism of action. The approach described here provides a direct way to screen for drug targets that would do this.

## Acknowledgements

This work was supported by the National Institutes of Health (R01-DK106399, R01-DK138127, and R01-DK145100 to Z.A.K.). Z.A.K. is an Investigator of the Howard Hughes Medical Institute. This work was also supported by the Postdoctoral Independent Research Grant (to S.Y.) co-funded by Program for Breakthrough Biomedical Research (partially funded by the Sandler Foundation) and UCSF EaRTH Center by P30-ES030284. Sequencing was partly performed at the UCSF CAT, supported by UCSF PBBR, RRP IMIA, and NIH 1S10OD028511-01 grants. Flow cytometry and cell sorting was performed at the UCSF LCA core facility, supported by P30CA082103. We thank Mark Moasser for access to the Seahorse Metabolic Analyzer. We thank Kivanc Birsoy and Denis Titov for helpful discussions at an early stage of this work.

## Methods

### Mouse lines

All experimental protocols using mice were approved by the University of California, San Francisco Institutional Animal Care and Use Committee, following the NIH Guide for the Care and Use of Laboratory Animals. CCK^Cre/+^ (#012706) and Igs7^GCamp8s/+^ (#037719) mice were obtained from Jackson labs.

### Molecular cloning

sgRNA-expressing constructs in pCRISPRia_v2: pCRISPRia_v2 vectors were digested with BstXI and BlpI and gel purified. DNA oligos with sgRNA sequences plus the overhang ends complementary to BstXI or BlpI sites were synthesized by IDT, phosphorylated by T4 PNK, annealed, ligated with digested pCRISPRia_v2 vectors and transformed in Stellar Competent Cells. All constructs were confirmed by Sanger sequencing.

### Lentivirus packaging and transduction

HEK293T cells were plated in 10-cm or 15-cm dishes so that they reach 40-50% confluency the next day. Transfection mix was prepared as following: transfer plasmid was mixed with psPAX2 and pCMV-VSVG at a 1:1:1 ratio, in Opti-MEM. 1 µg/µL polyethenylamine (PEI) was used at 3:1 ratio of total DNA mixture. DNA and PEI was then mixed and incubated at room temperature for 10 min and then added to HEK293T cells in a dropwise manner. Cell medium was collected each day for the following two days. The combined supernatant was passed through a syringe with a 0.45 µm filter. Lenti-X Concentrator was added at 1/3 volume of the filtered solution, which was mixed well and stored at 4°C for at least 4 h. Afterwards, the solution was centrifuged at 4°C for 45 min at 1,500 x g and the supernatant was discarded. The pellet was resuspended in cold media and aliquoted and stored at -80°C for future use.

Lentiviral transduction was performed with aliquoted lentivirus in complete media, with 8 µg/mL polybrene. The next day, the media containing lentivirus was discarded and cells were replaced with fresh media. Two days following transduction, the cells were dissociated and replated with desired selection drugs in the media.

### Generation of stable cell lines

Stable cell lines are generated by lentiviral transduction of desired constructs, followed by antibiotic selection for 1 week or sorting based on fluorescence. Single clones were then isolated by serial dilution or FACS-based plating. For ZIM3-KRAB-dCas9, the clone with the best knockdown efficiency was selected. For CaMPARI and GCaMP6m, the clones with best calcium responses were selected.

### CaMPARI photoconversion assay

Cells stably expressing CaMPARI were plated in 48-well plates the day before. Cells were washed briefly with DPBS and equilibrated in imaging buffer (20 mM HEPES, 140 mM NaCl, 2.5 mM KCl, 1.8 mM CaCl2, 1.0 mM MgCl2, pH 7.4) for 15 min. Cells were then treated with desired stimulus and photoconverted with UV (Formlabs, Form Cure, FH-CU-01) for 5 min. Cells were then rinsed with DPBS, dissociated with phenol red free TrypLE for 3 min, resuspended in FACS buffer (phenol red free DMEM, 2% FBS, 1% P/S) and plated in 96-well plates for flow cytometry analysis using Attune NxT equipped with an autosampler. The following channels were used: BL1 (488 nm excitation, 530/30 nm emission) for CaMPARI(green), YL1 (561 nm excitation, 585/16 nm emission) for CaMPARI (red), and VL1 (405 nm excitation, 450/40 nm emission) for BFP (when validating individual hits using sgRNA). 10,000 cells were collected for each sample. Flow cytometry data was analyzed using FlowJo (v10.10). Cells were gated with the FSC-SSC plot and intact singlets were then gated for CaMPARI (green) and BFP (sgRNA for CRISPRi KD samples). Red to green ratio was then calculated for each cell, and the median value for each sample was exported and plotted in Prism or R.

### Generation of a custom sgRNA library

Target genes in the custom sgRNA library include: 1) transmembrane signaling receptor activity (GO:0004888); 2) transmembrane transporter actvity (GO:0022857); 3) metabolic genes (amino acid metabolic process [GO: 0006520], electron transport chain [GO:0022900], ADP metabolic process [GO:0046031], and generation of precursor metabolites and energy [GO:0006091]); 4) hits from pilot screens done with Weissman m1 sublibrary; 5) top 150 differentially expressed genes in STC-1 compared to NIH3T3. After removing redundant genes and genes not expressed in STC-1 cells, top 5 predicted sgRNAs per gene (3744 genes and 19760 sgRNAs) plus 490 non-targeting sgRNA controls (for a total of 20250 elements) were synthesized by Twist and cloned into the pCRISPRia_v2 vector. Library complexity and element distribution was confirmed with Illumina next-generation sequencing by Twist and us independently.

### Pooled CRISPRi screening

Each FACS screen is paired with a survival screen using the same batch of cells and lentiviruses.

The sgRNA library was packaged into lentiviruses as described above. Functional viral titer was determined by measuring the BFP fluorescence using FACS following a small-scale test infection. STC-1 cells stably expressing dCas9 and CaMPARI were infected with the sgRNA library at MOIs between 0.3-0.5 with >1000x coverage per library element. Two days later, cells were selected with 10 µg/mL puromycin for two days. Then on Day 0, cells were cultured in complete media. A total number of cells >1000x coverage per library element were collected as D0 sample for survival screen. Cells were then maintained at a total cell number >2000x coverage per library element throughout. After cells underwent > 6x doubling, cells with a total number of >1000x coverage per library element were collected for survival screen. Additionally, cells were plated in 15-cm dishes for FACS sorting the next day. On FACS day, cells were treated with the desired stimulus and subject to CaMPARI photoconversion. Cells were dissociated, pelleted, resuspend in FACS buffer (phenol red free DMEM, 2% FBS, 1% P/S) and strained with 35 µm cell strainer. Top and bottom 35% of red-to-green ratio populations were sorted by BD FACSAria Fusion Flow Cytometer. Cell pellets were washed with PBS once and stored at -80°C until sample preparation.

For each screening sample (both survival screens and FACS screens), genomic DNA was isolated using Nucleospin Blood L Kit (Macherey-Nagel, 740954.20). Variable sgRNA regions were amplified with unique barcodes for each sample and sequenced on an Illumina HiSeq4000, NovaSeq6000 or NextSeq2000 as previously described (Gilbert et al., 2014; Kampmann et al., 2014).

Screens were analyzed as reported previously (Boggess et al., 2024). Briefly, sequencing reads were analyzed for sgRNA protospacer sequence and mapped to a reference library to generate a count matrix for each screen sample, using two publicly available Rust packages. Log2(fold change) and significance P values were calculated for target genes over “negative-control-quasi-genes”, as well as an FDR using the iNC method. Hits were defined with FDR < 0.05.

### qPCR

Cell pellets were thawed on ice, and total RNA was extracted using RNeasy Mini Kit (Qiagen, 74104). cDNA was synthesized using iScript cDNA Synthesis Kit (BioRad, 1708891). Samples were prepared for qPCR in triplicates, using PrimeTime qPCR probe assays (IDT; sequences in Supplementary Table x) in iTaq Universal Probes Supermix (BioRad, 1725134). Quantitative real-time PCR was performed on CFX Connect Real-Time PCR Detection System (Bio-Rad). Data was analyzed using the ΔΔCt method and normalized to *Gapdh*.

### Live cell calcium imaging

STC-1 or primary EECs were cultured in glass or polymer bottom black plates. Prior to imaging, cell medium was changed with imaging buffer and equilibrated for at least 10 min. Imaging was performed on a Nikon Ti2 widefield microscope with an Apochromat Lambda D 20x objective lens (FOV 25 mm) and a Digital Sight 50M camera.

Images were motion and background corrected, and cells were segmentated with Cellpose. ΔF/F0 was calculated as (F-F0)/F0, where F is the fluorescence intensity and F0 is the averaged baseline before stimulation. For primary EEC imaging, normalized ΔF/F0 was calculated as the ratio of tryptone response over KCl response.

### Bulk RNA-Seq

STC-1 and NIH3T3 cells were dissociated, pelleted and snap frozen as triplicates. Total RNA was extracted using the RNeasy Mini Kit (Qiagen, 74104). 3’ mRNA-Seq libraries were then prepared using the QuantSeq 3’mRNA-Seq Library Prep Kit (FWD) for Illumina (Lexogen, 015.24 & 020.96). Concentrations were measured with the Qubit 1X dsDNA HS Assay Kit on a Qubit Fluorometer. Library fragment-length distribution was quantified with the High Sensitivity D1000 ScreenTape on a TapeStation 4200 (Agilent). All libraries were diluted to equimolar representation in a pooled sample and submitted for single-end sequencing on an Illumina HiSeq4000.

After passing QC, sequencing reads were mapped to the *Mus musculus* reference transcriptome (GRCm39.cdna.all.release-105) and quantified by Salmon (v1.6.0). The Bioconductor package ‘tximetà(1.12.4) was used to convert transcript-level abundance to gene-level counts. Genes that have no counts or only a single count across all samples were removed. ‘DESeq2’(v1.34.0) was used to perform count normalization and differential expression analysis (with FDR threshold set to 0.05). When normalized counts were plotted on the log scale, a pseudo-count of 0.1 was added to all values.

### ELISA

Cells were plated in 24-well plates the day before. Cells were under treatment in imaging buffer for 2 h. Cell media were collected and centrifuged at 300 x g for 5 min and supernatant was collected. GLP-1 Total ELISA Kit (Sigma, EZGLP1T-36K or EZGLP1T-36BK) was used to measure GLP-1 concentration on BioTek Synergy LX multi-mode microplate reader.

### Primary EEC isolation and culture

Adult mice were euthanized, and proximal small intestine was immediately collected, chopped into around 3 cm sections, flushed with cold DPBS, opened longitudinally and rinsed thoroughly with DPBS. Each section was placed on a coverglass and examined under epifluorescence microscope to confirm expression of GCaMP in EECs. Villi were scraped off with coverglass.

Processed intestinal tissues were incubated in Dissociation Media I (DPBS, 30 mM EDTA, 10 µM Y-27632) on ice for 12 min, with brisk inversion every 2 minutes to dissolve mucous. Tissues were then transferred to prewarmed Dissociation Media II (DPBS, 30 mM EDTA) and incubated at 37°C for 6 min, with brisk shaking for 5 seconds every minute and after 6 min, vigorous shaking for 30-60 seconds or until epithelium was completed dissociated. Remaining large tissue pieces were removed and checked under a dissection microscope to confirm no epithelium remained attached. Subsequently, detached crypts were pelleted, washed, resuspended in prewarmed Digestion Buffer (HBSS, 0.1 U/mL Dispase II, 20 mM HEPES, 5 µM Y-27632, 100 µg/mL DNase I) and incubated at 37°C for 4-6 min, with vigorous shaking for 15 seconds every 30 seconds. Digested crypts were then pelleted, washed, strained with a 70 µm strainer, pelleted again and resuspended in complete media with B-27, 10 µM Y27632, and 250 µg/mL DNase I. The digestion resulted in a similar proportion of single cells and small clumps. Cells were plated on Matrigel-coated glass-bottom 24-well plates. After 2-3 hours, wells were flooded with media to remove unattached cells. Medium were changed daily to maintain cell survival.

### Seahorse Analyzer

Metabolic profiling with Seahorse Analyzer was performed following manufacture’s protocols on Agilent Seahorse XF 24, using the Seahorse XF Cell Mito Stress Test Kit. An acute injection cycle of DMSO or IACS010759 was added before the standard Oligomycin/FCCP/RAA cycles. Data were normalized with by cell counting (Hoechst staining) analyzed by Cytation.

---

**Supplemental Figure 1.**
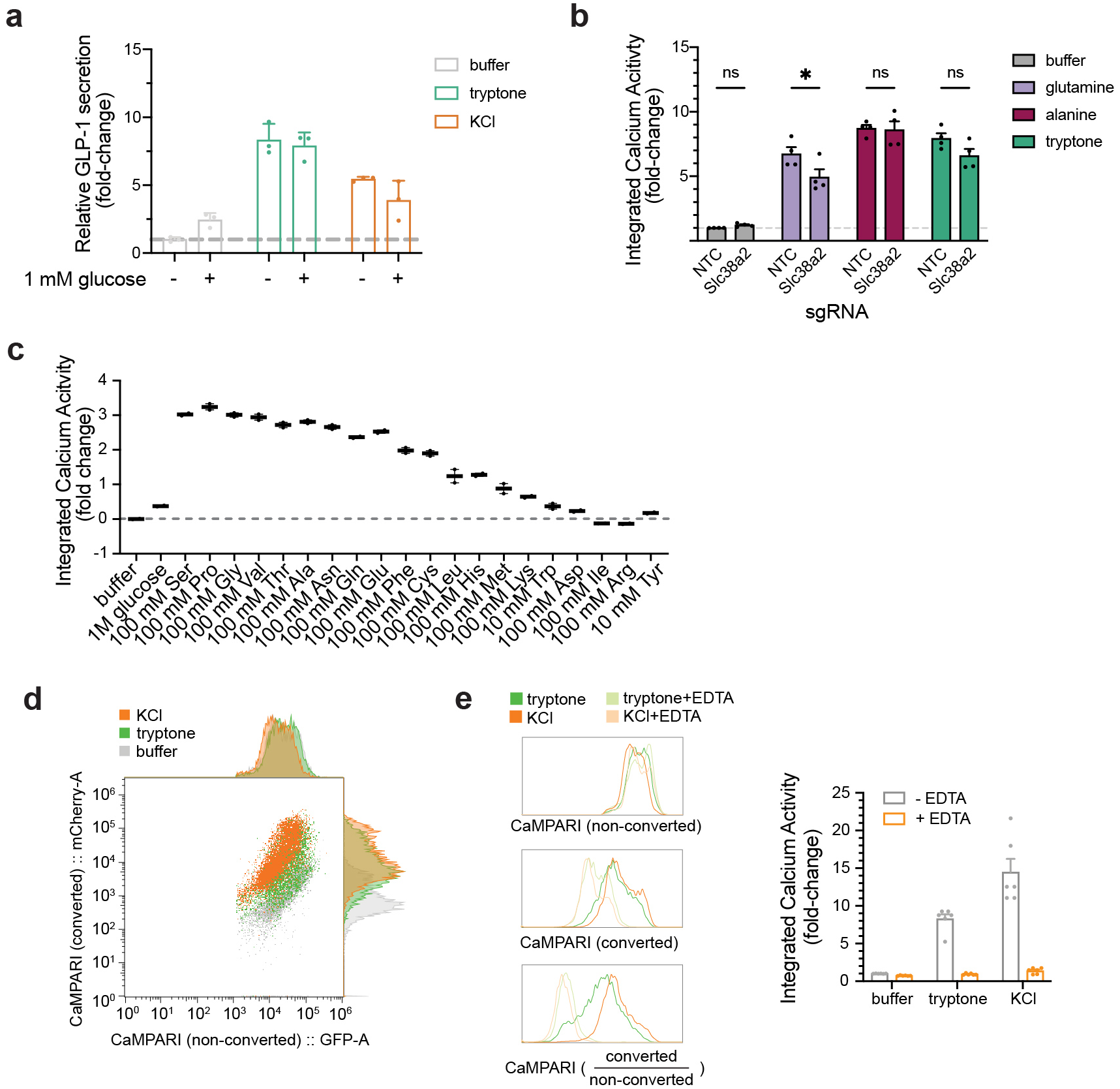
a, GLP-1 secretion in STC-1 does not depend on addtion of glucose. b, Integrated calcium activity for tryptone and Slc38a2 substrates after NTC or Slc38a2 KD. c, Integrated calcium activity (median CaMPARI photoconversion ratio) for individual amino acids in STC-1. d, Raw flow cytometry data for Figure 1h. e, EDTA completely inhibits stimulated calcium acitvity in STC-1. d, Histogram of CaMPARI activity under indicated conditions. e, Median CaMPARI photoconversion ratio from each flow cytometry session is plotted.

**Supplemental Figure 2.**
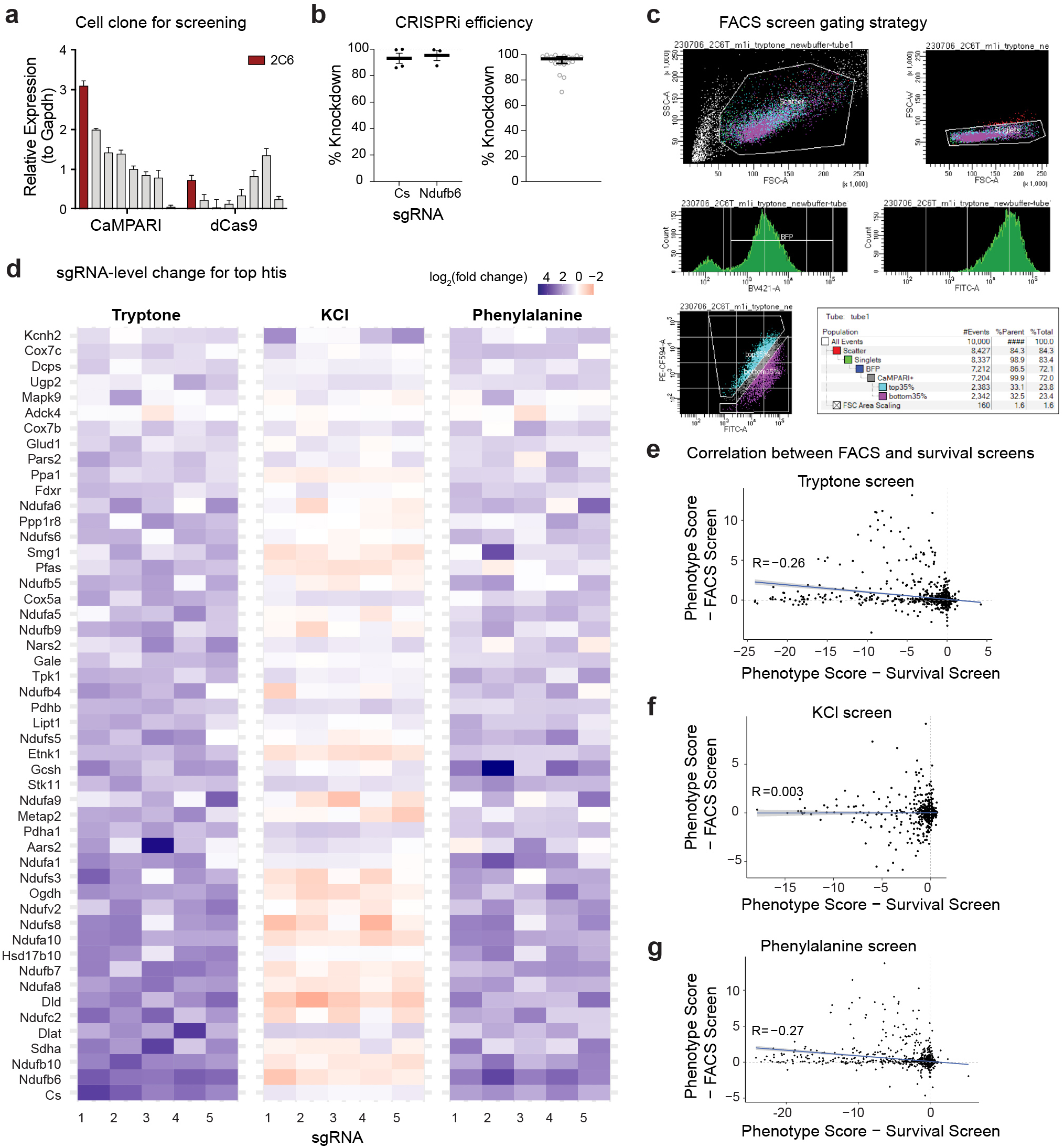
a, Expression levels of CaMPARI and ZIM3-KRAB-dCas9 in each isolated single clones. Clone 2C6 is selected for all the CaMPARI screens and validation shown in this manuscript. b, CRISPRi efficiency by qPCR. Left, Knockdown efficiency for two candidate genes shown in Figure 4. Right, Knockdown efficiency for all tested target genes. c, FACS screen gating strategy. Top and bottom 35% of CaMPARI photoconversion ratio (red/green) was collected. d, Heatmap showing log_2_(fold change) for each individual sgRNA (5 per gene) targeting the top 50 hits from the tryptone screen. e-g, Phenotype scores for all library genes, comparing FACS screen with interal survival screen control. e, Tryptone screen. f, KCI screen. g, Phenylalanine screen. Pearson correlation coefficient is shown on the plot.

**Supplemental Figure 3.**
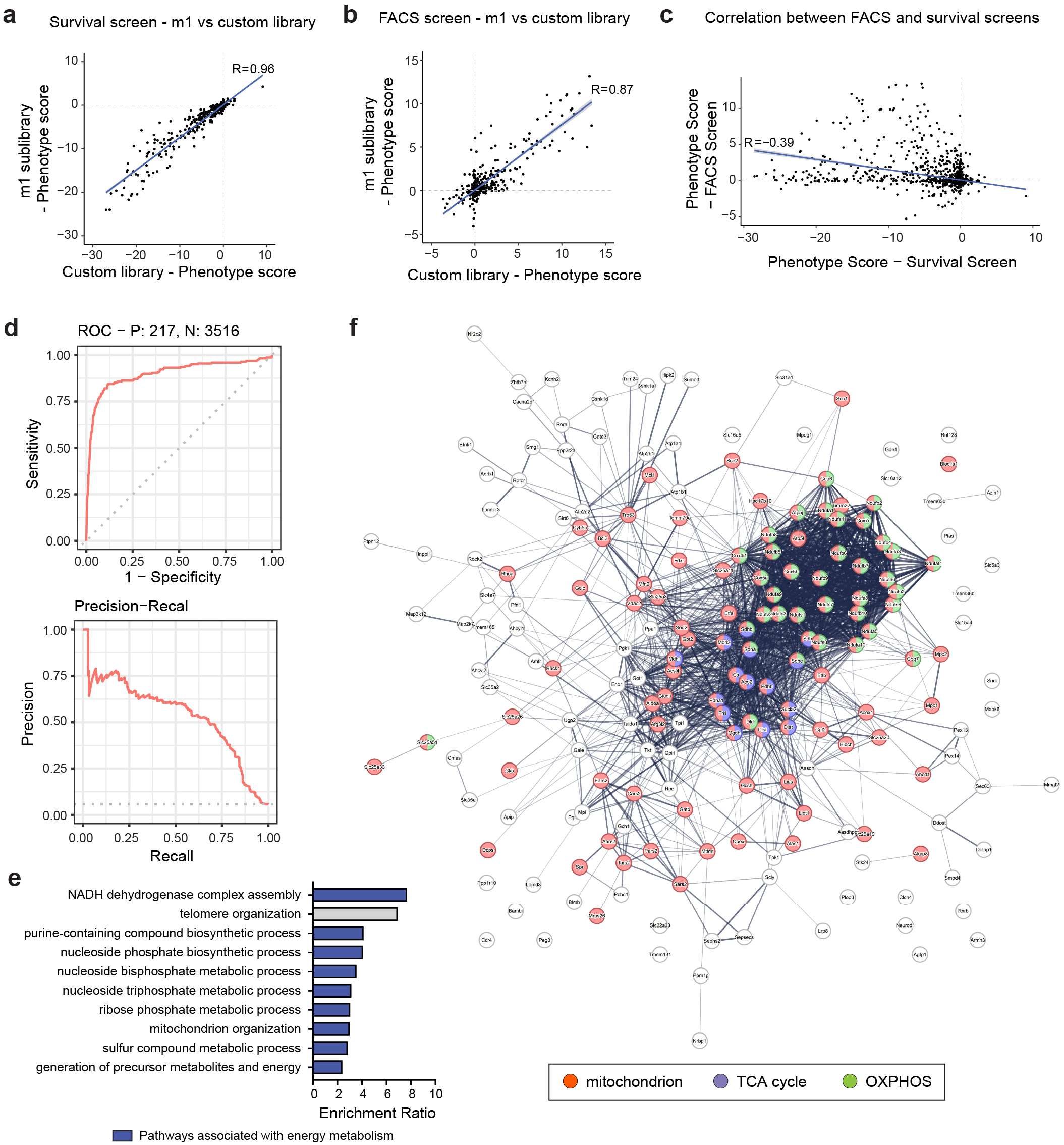
a-b, Correlation of phenotype score between m1 subpoole library screen vs. custom library screen. All hits present in both libraries are shown. a, Comparison between two survival screens. b, Comparison between two FACS screens. c, Correlation of phenotype score between FACS screen and survival screen for the large-scale custom library screen. d, Precision-recall analysis of survival screen showed reliable detection of essential genes. e, Top 10 most enriched pathways for all positive hits from the custom library screen. pathways associated with energy metabolism are highlighted. f, Functional protein-protein network analysis by STRING (as in Figgure 3d) labeled with all gene names. Genes involved in TCA cycle (GO:0006099) and OXPHOS (GO:0006119) are also highlighted.

**Supplemental Figure 4.**
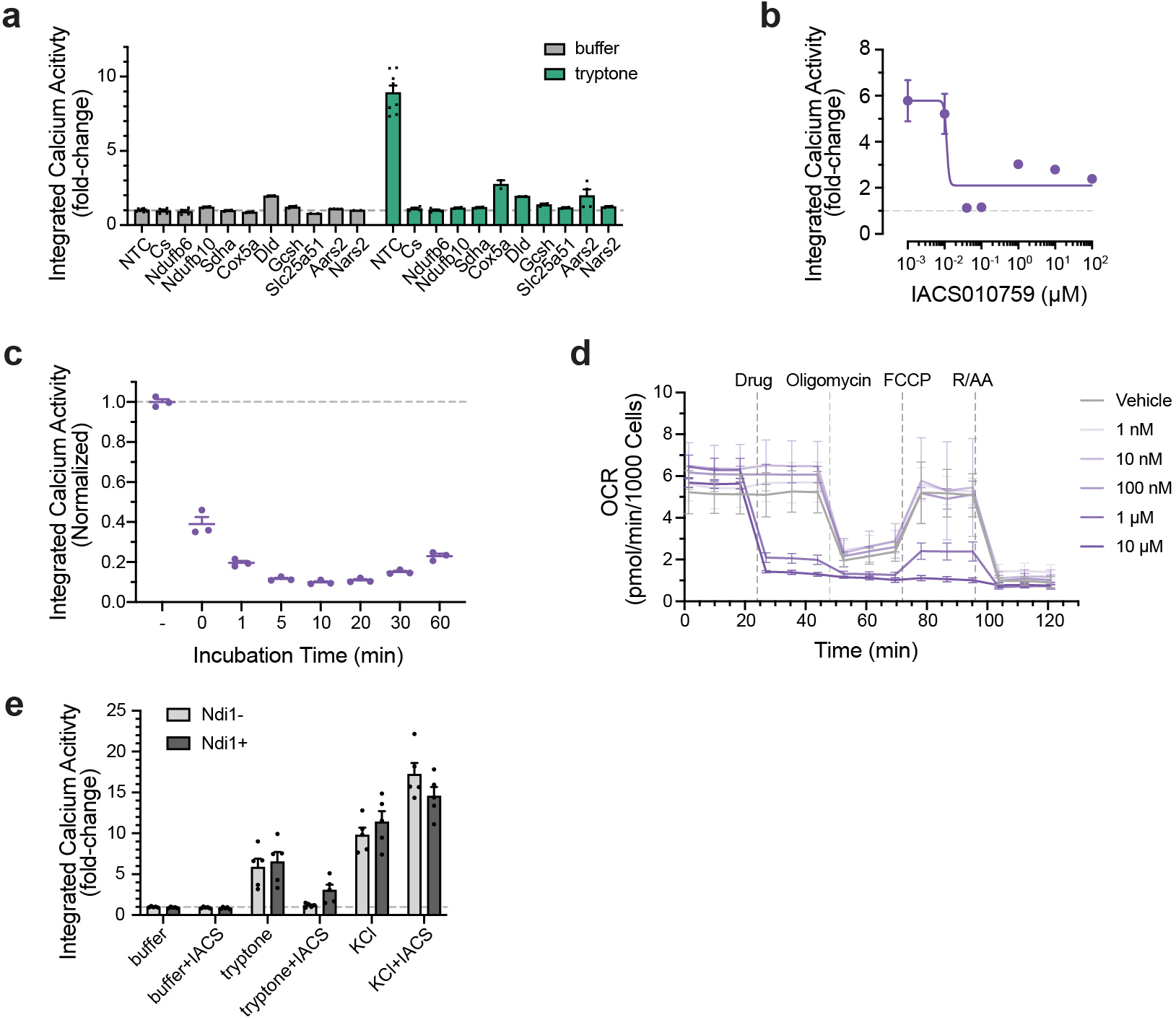
a, Validation of additional hits from the screen by CaMPARI. b, Dose response curve for the inhibitory effect of IACS010759 on tryptone-induced calcium activity. c, Inhibition of tryptone-induced calcium activity by IACS010759 after pretreatment of increasing length of time. d, Oxygen comsuption rate after vehicle or different concentrations of IACS010759 measured by Seahorse Analyzer. e, Integrated calcium activity with or without stable expression of yeast Ndi, the OXPHOS complex I equivalent. *P < 0.05; **P < 0.01; ***P < 0.001; ****P < 0.0001.

**Supplemental Figure 5.**
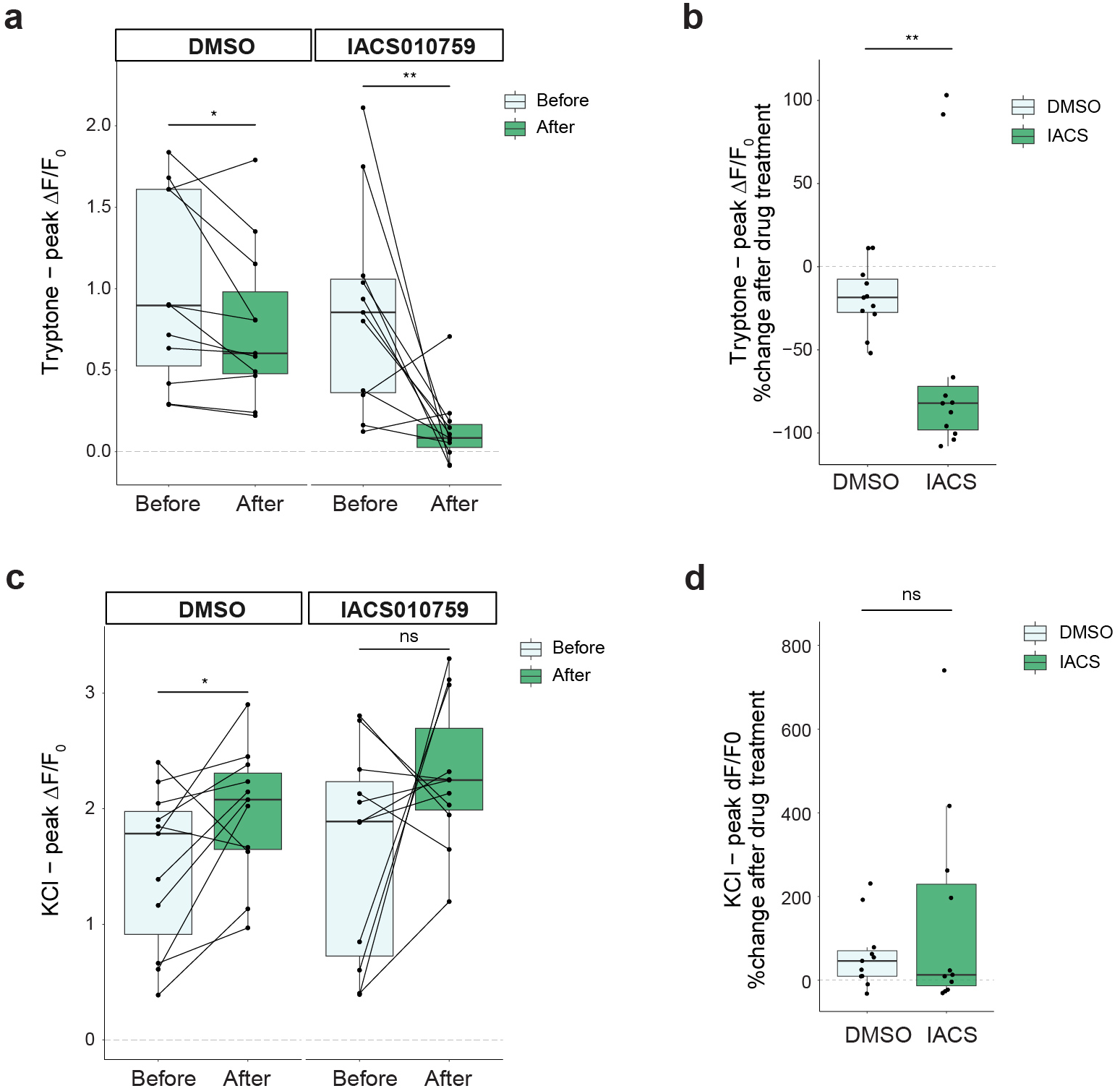
a-b, Quantification of calcium activity in primary EECs before and after treatment. Pairwise comparison of same cells is shown. a, Peak ΔF/F_0_ for tryptone. b, Percent of change after trytpone treatment for peak. ΔF/F_0_. c-d, Quantification of calcium activity in primary EECs before and after treatment. Pairwise comparison of same cells is shown. c, Peak. ΔF/F_0_ for KCI. d, Percent of change after KCI treatment for peak. ΔF/F_0_. *P < 0.05; ***P < 0.001; ****P < 0.0001.

